# vH^+^-ATPase-induced intracellular acidification is critical to glucose-stimulated insulin secretion

**DOI:** 10.1101/732222

**Authors:** Akshata R. Naik, Brent J. Formosa, Rishika G. Pulvender, Asiri G. Liyanaarachchi, Bhanu P. Jena

## Abstract

Swelling of secretory vesicles is critical for the regulated expulsion of intra-vesicular contents from cells during secretion. At the secretory vesicle membrane of the exocrine pancreas and neurons, GTP-binding G proteins, vH^+^-ATPase, potassium channels and AQP water channels, are among the players implicated in vesicle volume regulation. Here we report in insulin secreting MIN6 cells, the requirement of vH^+^-ATPase-mediated intracellular acidification on glucose-stimulated insulin release. MIN6 cells exposed to the vH^+^-ATPase inhibitor Bafilomycin A show decreased acidification of the cytosolic compartment that include insulin-carrying granules. Additionally, a loss of insulin granule association with the cell plasma membrane is demonstrated following Bafilomycin A treatment and results in a decrease in glucose-stimulated insulin secretion and accumulation of intracellular insulin. These results suggest that vH^+^-ATPase-mediated intracellular acidification is required both at the level of secretory vesicles and the cell plasma membrane for cell secretion.

## INTRODUCTION

Osmotic swelling of intracellular vesicles was first reported in sea urchin eggs^1,2^. Studies in mast cells show that osmotic swelling of secretory vesicles is important for their fusion with the cell plasma membrane and in fusion pore dilation^3^. Thereafter, the requirement of vesicle swelling in exocytosis in different tissues including neurons^4,5^ have been demonstrated. The extent of vesicle swelling following a secretory stimulus has been found to be directly proportional to the amount intra-vesicular contents released from the cell^5^. The molecular mechanisms underlying the swelling process has also been greatly explored and elucidated. Aquaporins (AQP), the rapidly gating water channels are present at the secretory vesicle membrane^6-8^ and participate in vesicle volume regulation. At least 13 different aquaporin isoforms have been identified. Among them, AQP1 has been demonstrated to be present at the membrane of zymogen granules, the secretory vesicles in the exocrine pancreas carrying digestive enzymes. Similarly, AQP6 has been found to be present at the synaptic vesicle membrane in neurons. Studies further report aquaporins to be under the regulation of GTP-binding G proteins, such as G_αi_3 in zymogen granules^4-6^ and G_αo_ in the neuronal synaptic vesicles^9^. Studies suggest that the electrochemical gradient generated by the influx of protons and potassium, are among the major driving force for AQP-mediated water entry into secretory vesicles^4-6,9^. The proton pump, vacuolar vH+-ATPase present at the synaptic vesicle membrane operates upstream of G_αo_-induced aquaporin mediated synaptic vesicle swelling^9^.

In insulin-containing granules of the endocrine pancreas, G_αi_ is localized to the granule membrane and stimulates insulin secretion^10^. Furthermore, acidic pH of insulin-containing intragranular lumen has been demonstrated to be important for the conversion of insulin from its prohormone to its mature form in the granule^11-13^. Based on these earlier reports, we hypothesized that insulin secreting beta cells may require intragranular acidification similar to zymogen granules of the exocrine pancreas and neurons, enabling vesicle swelling and insulin secretion. In the current study, to test this hypothesis, we used MIN6 mouse insulinoma cells and subjected them to Bafilomycin A, a pharmacological inhibitor of vH^+^-ATPase to inhibit cellular and insulin secreting granule acidification. We found significant loss in potency and efficacy of glucose-stimulated insulin release in Bafilomycin A-treated MIN6 cells compared to unexposed controls. Additionally, the accumulation of insulin within Bafilomycin A treated MIN6 cells was demonstrated.

## EXPERIMENTAL PROCEDURES

### Estimation of glucose-stimulated insulin secretion from cultured MIN6 cells

MIN6 cells were grown to confluency using sterile 100 × 13-mm plastic Petri dishes according to published procedure^14^. Cells were cultured in 25 mM glucose Dulbecco’s Modified Eagle Medium (Invitrogen, Carlsbad, CA) containing 10% fetal calf serum, penicillin, streptomycin, and 50μM *β*-mercaptoethanol. Cells were stimulated using 35 mM glucose and insulin secreted into the medium was collected at 0 min, 10 min and 30 min post stimulation, for determination of insulin using Western blot analysis. Stimulation assays were performed on MIN6 cells pre-exposed for 10min to 0, 10 and 50 nM Bafilomycin A. All secretion assays were performed at room temperature (RT) on cells prewashed in phosphate buffered saline (PBS) pH 7.4 followed by incubation in PBS. Following glucose stimulation, 200 μL aliquots of the incubation medium were collected at 0 min, 10 min and 30 min post stimulation. Aliquots were centrifuged at 4,000 *xg* to separate any aspirated cells. 160 μL of the cleared supernatant was mixed with 40 μL of 5x Laemmli reducing sample preparation buffer^15^ for Western blot analysis. To obtain the total amount of insulin in cells, Min6 cells were thoroughly scraped from the culture plates and subjected to solubilization in 100 μL of homogenization butter (2mM EDTA, 2mM ATP, 0.02% Triton X-100, 1:500 protease inhibitor cocktail, pH 7.4), following secretion assays. Protein in the solubilized cell fractions were determined using the Bradford protein assay^16^ prior to Western blot analysis.

### Detection of intracellular pH change

Changes in intracellular pH was determined using acridine orange (AO). AO is a membrane penetrating dye which is a weak base that accumulates within acidic cellular compartments. AO binds to double stranded DNA emitting green fluorescence and to single stranded DNA or RNA emitting red fluorescence. Other acidic compartments such as the lysosome and secretory vesicles in the red and green merged images fluoresce orange. AO emits a bright orange fluorescence at lower pH, turning to lighter orange at relatively basic pH. MIN6 cells were cultured on 35mm glass bottom petri dishes and treated with either 10nM Bafilomycin A, 50nM Bafilomycin A or vehicle. To determine the relative acidity of various cellular compartments, MIN6 cells were incubated with AO (2 μg/ mL) for 20 minutes at 37° C following treatment with either Bafilomycin A or vehicle and washed with sterile 1x PBS, pH 7.4. Following fluorescence microscopy, orange acidic compartments in the red and green merged images were observed.

### Western blot analysis

Min6 cell lysates (10 μg) and 20 μL of the cell incubation medium in Laemmli buffer, were resolved using 12.5% SDS-PAGE and electro-transferred to 0.2mm nitrocellulose membrane for immunoblotting and detection using chemiluminescence. The nitrocellulose membrane was incubated at RT for 1h in blocking buffer (5% non-fat milk in PBS-0.1% Tween pH 7.4), washed thrice with PBS pH 7.4 containing 0.1% Tween, and immunoblotted at 4°C overnight with mouse polyclonal monospecific anti-insulin antibody (2D11-H5) (SC 8033). Prior to overnight incubation at 4°C with secondary antibodies (Donkey anti-Rabbit Alexafluor 594 (Invitrogen A21207), the nitrocellulose membranes were washed in PBS-0.1% Tween pH 7.4, thrice. Immunoblots were processed for enhanced chemiluminescence, exposed to X-Omat-AR film, developed and analyzed using ImageJ.

### Immunocytochemistry

MIN6 cells were grown on 35-mm glass bottom Petri dishes for immunocytochemistry. The distribution of anti-Gα_i3_ and insulin containing granules in 10 nM and 50 nM Bafilomycin A-treated (10min at 37°C) Min6 cells were compared with vehicle treated control cells. Primary antibodies: rabbit polyclonal anti-Gα_i3_ (C-10, SC 262) and mouse polyclonal anti-insulin (2D11-H5, SC 8033) and secondary antibodies: Donkey anti-Rabbit Alexafluor 594 (Invitrogen A21207) and donkey anti-Mouse Alexafluor 488 (Life technologies A21202), were used in the study. Cells were exposed to 4′,6-diamidino-2-phenylindole (DAPI) nuclear blue stain in 1X PBS (0.2µM) for nucleus localization. An immunofluorescence FSX100 Olympus microscope was used to acquire immunofluorescent images through a 63x objective lens (numerical aperture, 1.40) with illumination at 405, 488, or 647nn. Insulin and Gα_i3_ localization and cellular distribution were obtained through merging fluorescent images using ImageJ.

## RESULTS AND DISCUSSION

### Bafilomycin A inhibits acidification of the cytosolic compartment containing insulin secreting granules

Bafilomycin A and concanamycins are a related family of pleicomacrolide antibiotics derived from the Streptomycetes species. They are highly specific pharmacological inhibitors of vH^+^-ATPase without affecting any other type of ATPase pump^17,18^. MIN6 cells were incubated with acridine orange, which accumulated within intracellular acidic compartments. Acridine orange binds to double stranded DNA and exhibits emission maxima at 525 nm (Green) and when bound to single stranded DNA or RNA, the emission maxima is at 650 nm (Red). Besides the nucleus, other acidic compartments in cells such as the lysosome and secretory vesicles are known to stain with acridine orange. While the red and green staining of the nucleus is unaffected by exposure of MIN6 cells to the vH^+^-ATPase inhibitor Bafilomycin A [Figure 1], there is a clear decrease in staining of the cytosolic compartment where insulin secreting granules are located in Bafilomycin A-treated cells [Figure 1]. Bright orange fluorescence in the merged red and green images is observed in control MIN6 cells depicting acidified cytosolic compartment of cells where insul secretory granules are present. However, MIN6 cells pretreated with 50 nM Bafilomycin A exhibited a quenched orange fluorescence, suggesting secretory vesicle alkalization due to the inhibition of vH^+^-ATPase activity at the cell plasma membrane and at the membrane of insulin containing granules.

**Figure 1:**
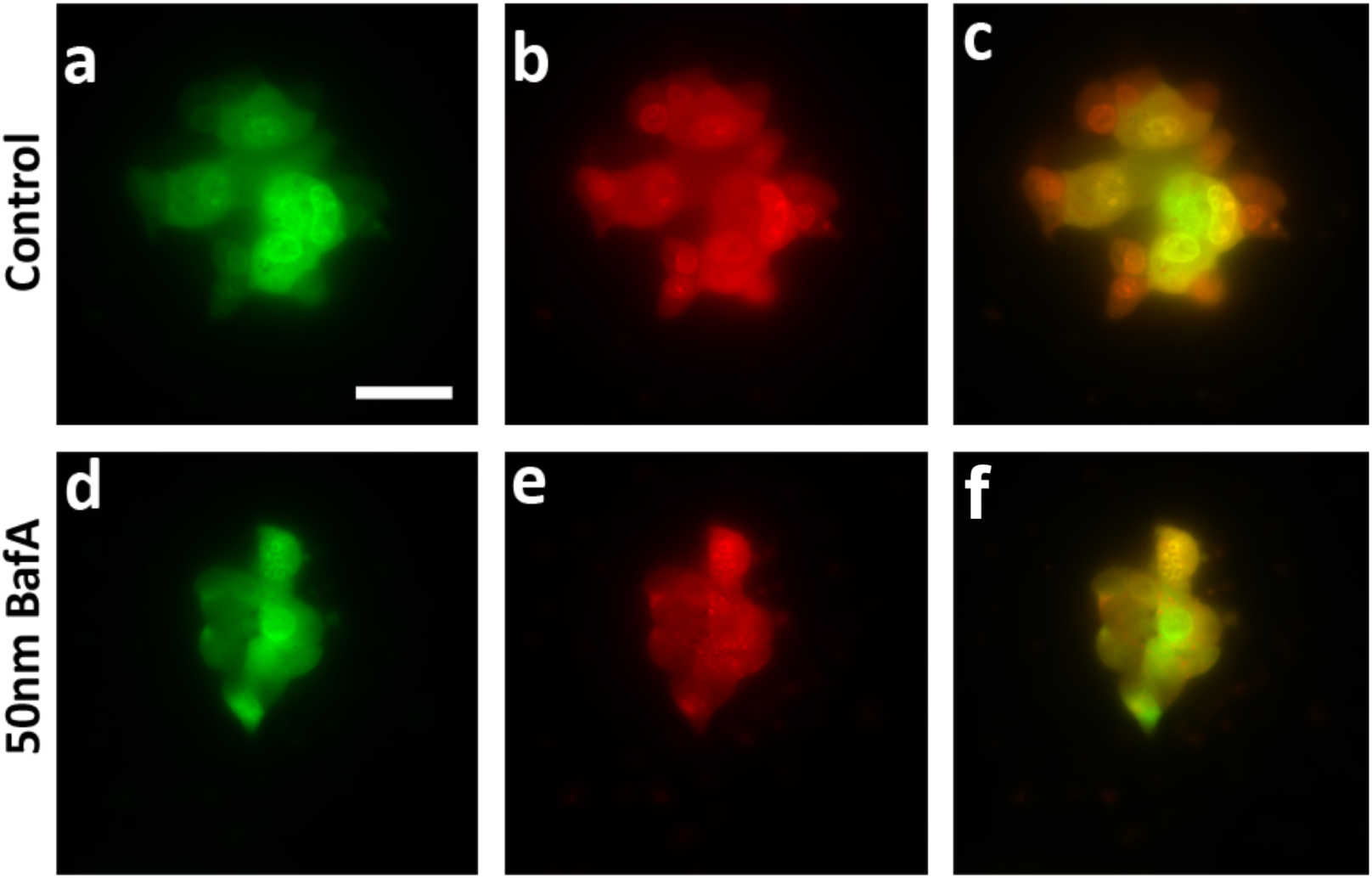
Bafilomycin A inhibits cytosolic acidification of MIN6 cells. MIN6 cells were incubated with 50 nM Bafilomycin A for 10 minutes at 37°C and loaded with acridine orange. Note the red and green staining of the nucleus is unaffected by exposure of MIN6 cells to the vH^+^-ATPase inhibitor Bafilomycin A. However, there is a visible decrease in staining of the cytosol where insulin secreting granules are located in Bafilomycin A-treated cells. This is further illustrated by the drop in bright orange fluorescence in the merged images in the Bafilomycin A-treated (f) cells compared to controls (c). *Scale bar =* 50 μm.

### Bafilomysin A-exposed MIN6 cells exhibit loss of glucose-stimulated insulin secretion

The importance of pH in secretion^4,5,6,9^ and granule maturation^19^ is well established. Therefore, our objective was to test whether Bafilomycin A treatment resulting in the inhibition of proton transport into cellular compartments including insulin-containing secretory vesicles, impacted glucose stimulated insulin release. We observed a significant reduction (p< 0.05) in glucose-stimulated insulin secretion upon Bafilomycin A treatment of MIN6 cells [Figure 2]. At both 10 nM and 50 nM Bafilomycin A concentrations, MIN6 cells exhibit significantly lower insulin release compared to vehicle-treated controls. Both the potency and efficacy of glucose-stimulated insulin release was reduced in Bafilomycin A treated MIN6 cells [Figure 2].

**Figure 2:**
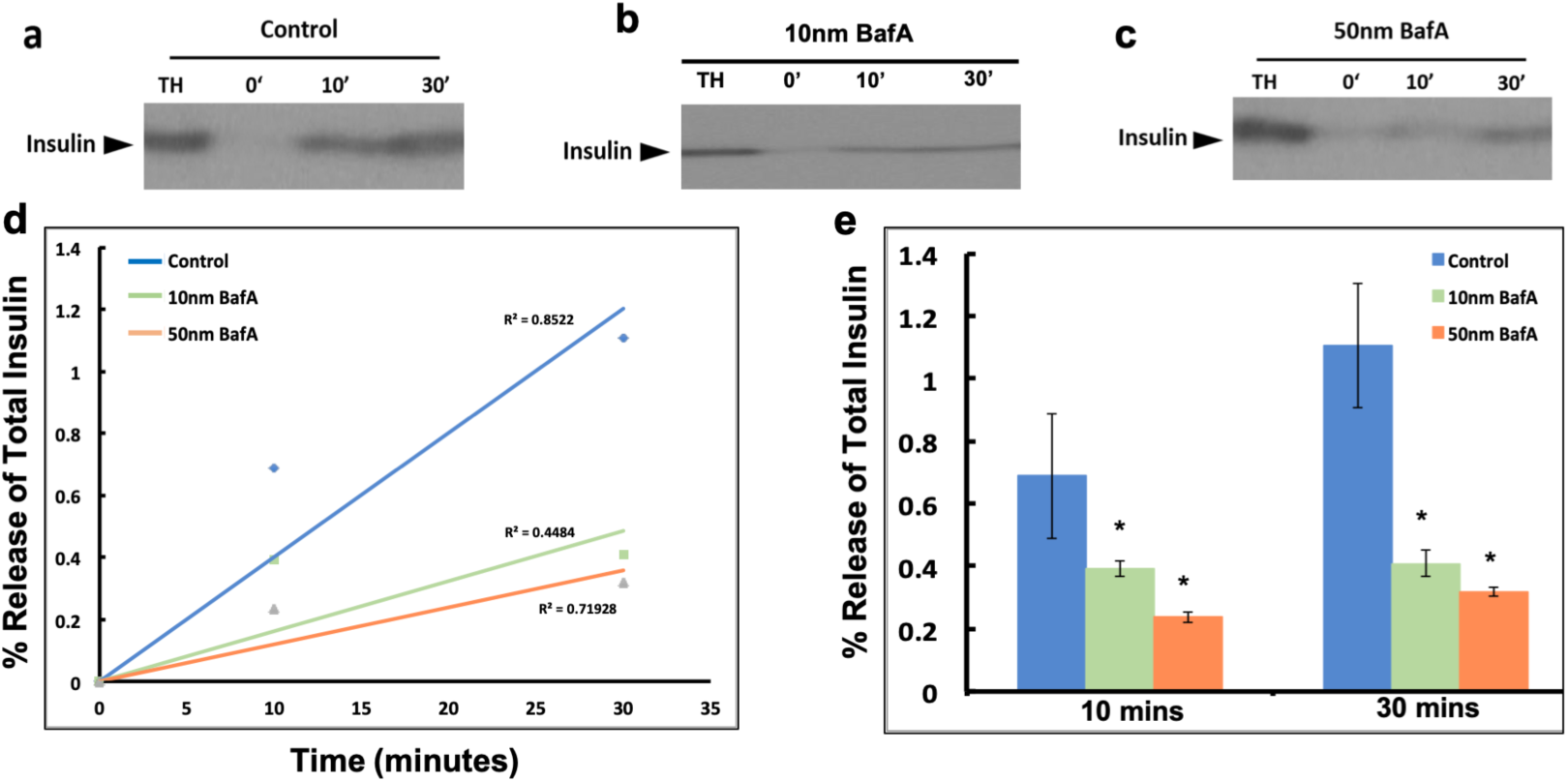
Bafilomycin A treatment reduces glucose-stimulated insulin release in MIN6 cells. Western blots on total homogenate (TH) from control MIN6 cell and cell medium at 0, 10 and 30 min post glucose stimulation (a), compared to 10 nM (b) and 50 nM (c) Bafilomycin A (BafA) treatment. Significant decrease (*p< 0.05; n=3 separate experiments) in insulin release is demonstrated at 10 min and 30 min post glucose stimulation of MIN6 cells. (d,e) Time course plots of percent insulin release demonstrate a significant drop in both the rate and amount of glucose-stimulated insulin release following Bafilomycin A treatment.

### Exposure of MIN6 cells to Bafilomycin A reduces insulin granule association with the cell plasma membrane

Immunocytochemistry performed on control and Bafilomycin A-treated MIN6 cells demonstrate loss of association of insulin containing vesicle with the cell plasma membrane. Antibody against insulin used to determine the distribution of insulin secreting granules in MIN6 cells and similarly antibody against G_αi_3 used as a plasma membrane and porosome (secretory vesicle docking and fusion machinery at the cell plasma membrane) marker, was used to determine interactions between them following exposure of MIN6 cells to Bafilomycin A. Exposure to either 10 nM and 50 nM Bafilomycin A demonstrated cytosolic accumulation of insulin secreting granules and their reduced co-localization with G_αi_3 at the cell plasma membrane as opposed to control cells [Figure 3]. Furthermore, an increase in insulin labeling is observed in Bafilomycin A-treated cells compared to untreated controls, demonstrating intracellular insulin accumulation may be due to a decrease in ‘basal’ or so called ‘constitutive secretion’ [Figure 3].

**Figure 3:**
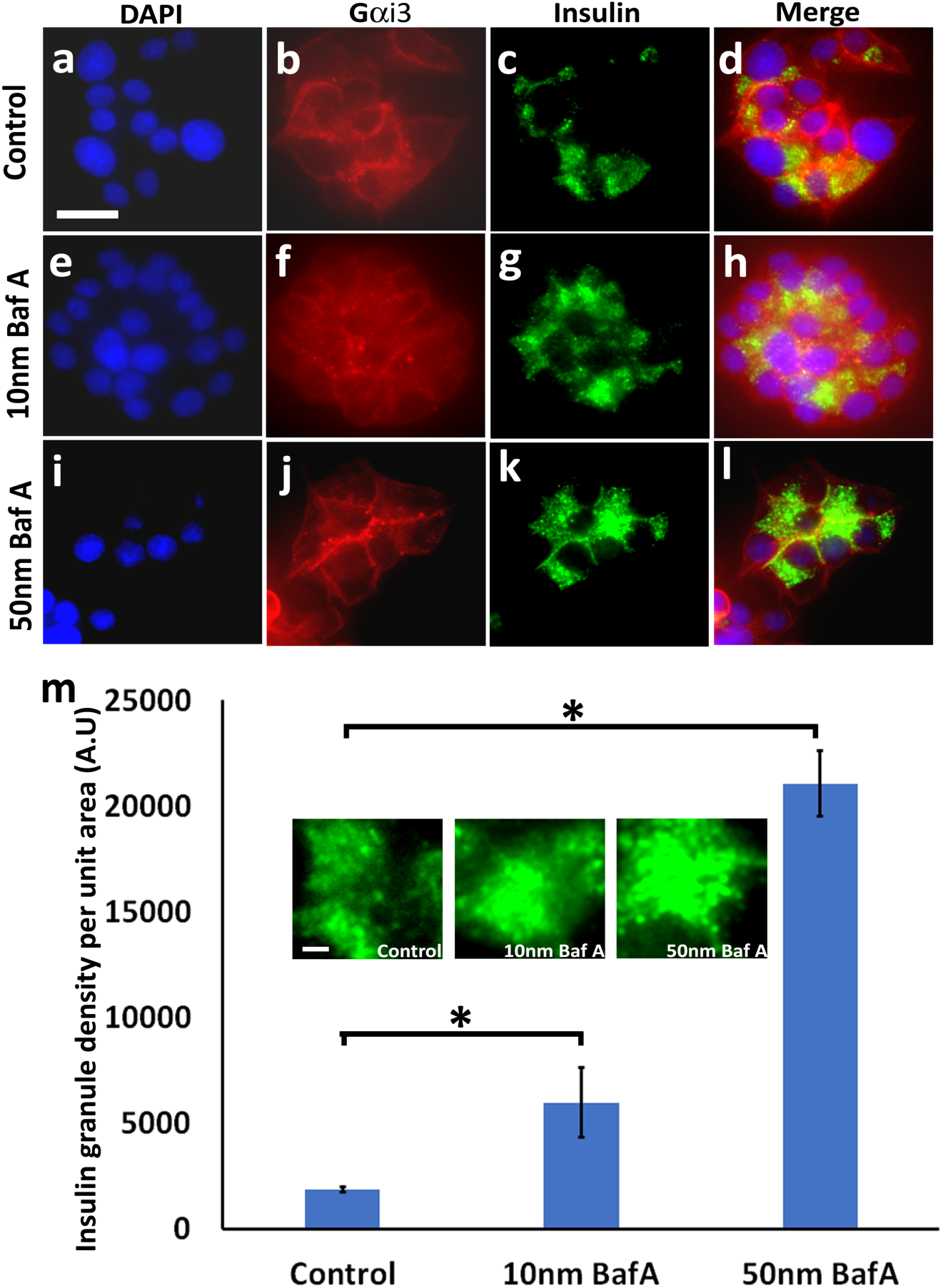
Bafilomycin treatment leads to accumulation of insulin within MIN6 cells. Fluorescent labeling of MIN6 cell nucleus using DAPI (Blue; a,e,i), immunofluorescent labeling of the cell plasma membrane using a Gαi-specific antibody followed by secondary rhodamine (Red; b,f,j) and immunofluorescent labeling of the insulin containing secretory granules using a insulin-specific antibody followed by a secondary green fluorescent antibody (Green; c,g,k), enables an understanding of the subcellular distribution of insulin containing granules. Note in the merged images, the increase in insulin granules in 10 nM Bafilomycin A-treated (h) and 50 nM Bafilomycin A-treated (l) cells compared to control cells (d). (*Scale bar = 100 μm*). ImageJ analysis demonstrates significantly (*p < 0.05) higher antigen density of insulin granules in Bafilomycin A-treated cells as compared to controls (m); Insets depict digitally zoomed regions of equal areas of the cytosol containing insulin labeling (green), suggesting increased insulin granules accumulation in both 10 nM and 50 nM Bafilomycin A treated cells over control.

Results from the study demonstrates that similar to synaptic vesicles in neurons and zymogen granules in the exocrine pancreas, insulin secreting granule acidification in MIN6 cells is critical for insulin release. When vesicle acidification is prevented by cellular exposure to Bafilomycin A to inhibit vH^+^-ATPase, insulin release dropped significantly. Similar to synaptic vesicles where Go, potassium ion channel, vH^+^-ATPase and AQP6 are among the players involved in volume regulation required for neurotransmitter release, insulin vesicle swelling as a consequence of rapid water gating via AQP7 previously identified in beta cells^20^ could be involved in glucose-stimulated insulin secretion. Furthermore, these results are in confirmation with recent studies^21^ demonstrating that Valproate, the FDA approved antiepileptic drug perturbs insulin granule-associated vH^+^-ATPase by preventing the association of the vH^+^-ATPase subunit C to the complex, thereby inhibits glucose-stimulated insulin secretion from MIN6 cells. Similarly, the critical role of intracellular acidification on decreasing t-/v-SNARE complex disassembly by the soluble *N*-ethylmaleimide-sensitive factor NSF, promoting fusion of secretory vesicles at the cell plasma membrane^22^, has also been demonstrated. Taken together, these results demonstrate the critical role of intracellular acidification on cell secretion both at the level of secretory vesicles and at the cell plasma membrane. The final step of the secretory process, i.e., the docking and fusion of secretory vesicles at the base of cup-shaped lipoprotein structures called porosomes^14,23-28^ in the cell plasma membrane. However, the molecular details of the exact sequence of events that follow insulin granule acidification are important and is the subject of current studies in the laboratory.

## Author Contributions

A.R.N and B.P.J designed the experiments and wrote the manuscript. B.F.J, R.G.P and A.L analyzed results and contributed to methods section. All authors critically analyzed results and proof read the manuscript.

## Acknowledgements

Work presented in this article was supported in part by WSU Interdisciplinary Biomedical Systems Fellowship.

## Competing Financial Interest

The authors declare no competing financial interest.

